# Divergent specificity of PatA, GabT, and IlvE defines the branched transamination of *N*_ε_-carboxymethyllysine and its metabolite *N*_ε_-carboxymethylcadaverine in *Escherichia coli*

**DOI:** 10.64898/2026.06.15.732455

**Authors:** Patroklos Vougioukas, Erica F. Aveta, Vincent Hoffmann, Jürgen Lassak, Michael Hellwig

## Abstract

Thermal food processing generates *N*_ε_-carboxymethyllysine (CML), a key advanced glycation end product (AGE) and marker of the Maillard reaction in food. *Escherichia coli* utilizes CML as a nitrogen source. While SpeC initiates degradation by decarboxylating CML to *N*-carboxymethylcadaverine (CM-Cad), the enzymes liberating the nitrogen remained unknown. Here, we identify PatA, GabT, and IlvE as the glutamate-dependent transaminases responsible for CML and CM-Cad transamination. Our results reveal a branched metabolic network rather than a linear pathway: PatA shows specificity towards both substrates, while GabT and IlvE selectively process CM-Cad and CML, respectively. We further demonstrate that the carboxymethyl piperideinium ion (CM-Pip) is formed spontaneously following CM-Cad transamination and reveal the previously unknown carboxymethyl-tetrahydropicolinic acid (CM-THPA) as novel metabolite in CML metabolism. Combining molecular microbiology, biochemistry, and analytical chemistry, we demonstrate that these transaminases are essential for integrating dietary CML into bacterial nitrogen metabolism, providing a model for microbial AGE processing via underground metabolism.

## 1. Introduction

Thermal food processing initiates the Maillard reaction, a complex network of non-enzymatic reactions between reducing sugars and amino groups (Hellwig et al., 2015; Hellwig et al., 2024; Lassak et al., 2023). This process begins with the formation of reversible Schiff bases, which subsequently rearrange into more stable early glycation products, among them Amadori products such as fructoselysine (Graf von Armansperg et al., 2021). Through successive rearrangements, dehydration, and fragmentation, these intermediates progress to form advanced glycation end products (AGEs) (Lassak et al., 2019; Lassak et al., 2022). Among these, *N*_ε_-carboxymethyllysine (CML) has long been established as a prominent and stable marker for monitoring the heat impact of food products (Hellwig et al., 2024). While its presence in the human diet and its role as a diagnostic marker have been known for decades, its interaction with the human gut microbiota has only recently gained scientific attention.

The capacity of intestinal bacteria to degrade CML was first described in detail in 2019. Here, we characterized the metabolization of CML by probiotic *E. coli* strains, identifying various carboxymethylated degradation products (Hellwig et al., 2019). Concurrently, a parallel study by Bui *et al*. identified anaerobic gut bacteria, such as *Oscillibacter spp.* and *Cloacibacillus evryensis*, as CML degraders, yielding similar carboxymethylated biogenic amines and cyclic derivatives (Bui et al., 2019). While these works established the general existence of a metabolic pathway and identified key intermediates such as CM-Cad and CM-Pip, the specific enzymes responsible for these transformations remained entirely unknown at that time.

To define the enzymatic basis for these metabolic observations, we recently identified the first enzyme of this pathway by demonstrating that the decarboxylation of CML to CM-Cad is mediated by the underground activity of the constitutive ornithine decarboxylase SpeC (Aveta et al., 2026). Underground activity, or enzyme promiscuity, refers to the capacity of enzymes to catalyze secondary reactions with non-native substrates that are structurally similar to their primary targets (D’Ari & Casadesús, 1998; Guzmán et al., 2015). This phenomenon is a cornerstone of metabolic flexibility and evolution, as it allows organisms to exploit novel environmental compounds by recruiting existing catalytic scaffolds for alternative functions (Richts & Commichau, 2021; Rosenberg & Commichau, 2019). In the case of CML, the recruitment of SpeC provides a dual physiological benefit: it supports the pH stress response via proton consumption and initiates the liberation of nitrogen (Aveta et al., 2026). Our previous research established that while *E. coli* can utilize CML as a weak nitrogen source, it cannot use it as a sole carbon source. Although the identification of SpeC resolved the first enzymatic step, the enzymes required to process the resulting variety of intermediates and fully integrate the nitrogen into central metabolism remained to be discovered.

In the present study, we identify the next enzymes in the CML degradation network of *E. coli*. We demonstrate that this process is not a simple linear pathway but a branched metabolic network where both CML and its decarboxylation product CM-Cad serve as substrates for transamination. We identified the glutamate-dependent transaminases PatA, GabT, and IlvE as the key enzymes in this process. Specifically, we show that PatA exhibits substrate specificity towards both CML and CM-Cad, while GabT and IlvE act more selectively, processing CM-Cad and CML, respectively.

Furthermore, we characterize the resulting metabolic products, including the spontaneously formed CM-piperideinium ion and carboxymethyl-tetrahydropicolinic acid (CM-THPA). By combining *in vitro* biochemistry, *in vivo* mutant analysis and analytical chemistry, we provide a comprehensive model for how dietary CML is integrated into the bacterial nitrogen metabolism via promiscuous transaminase activities.

## 2. Materials and methods

### 2.1 Media and growth conditions

*E. coli* was cultivated in either Lysogeny Broth (LB) (Bertani, 1951; Miller, 1972) at 37 °C, or M9 minimal medium (Green & Sambrook, 2012) (33.7 mM Na_2_HPO_4_, 22.0 mM K_2_HPO_4_, 8.55 mM NaCl, 10 mM NH_4_Cl, 1 mM MgSO_4_, 0.3 mM CaCl_2_, 1 μM biotin and 10 µM thiamin, 13.4 µM ethylenediaminetetraacetic acid (EDTA), 3.1 µM FeCl_3_, 0.62 µM ZnCl_2_, 76 nM CuCl_2_, 42 nM CoCl_2_, 162 nM H_3_BO_3_, 8.1 nM MnCl_2_), supplemented with 20 mM D-glucose or 20 mM glycerol as the carbon source and different nitrogen sources as indicated. If required, other media supplements or antibiotics were added to the culture. For induction of protein overexpression in *E. coli* LMG194 and BW25113, L-arabinose was added to the media in the logarithmic growth phase. For long-term storage of bacterial strains, cell cultures were snap-frozen in liquid nitrogen and stored at −80 °C in glycerol freezing medium (87% glycerol, 1 M KCl, 1 M NaCl, 0.2 M MgSO_4_) in a 1:3 ratio. Before inoculation in M9, the cells were washed twice (15,000 × *g*, 1.5 min) in M9 lacking carbon and nitrogen to prevent nutrient carryover. Growth was recorded by measuring optical density at 600 nm (OD_600_) using either a photometer (Ultrospec 2100 pro UV-Vis spectrophotometer, Amersham Biosciences, UK) or a plate reader spectrophotometer equipped with a thermostat (CLARIOstar Plus, BMG LABTECH, Germany). The strains used in this study are listed in Table S1.

### 2.2 Cloning and plasmid preparation

*E. coli* cells were prepared for electroporation by washing twice with ice-cold 10% glycerol, and subsequently transformed using a Bio-Rad MicroPulser (Bio-Rad, USA) set to the bacterial cell mode. The CDSs of the selected transaminase genes (*argD*, *aspC*, *astC*, *gabT*, *ilvE*, *patA*, *puuE*, *tyrB*, and *ybdL*) were amplified using Q5^®^ High-Fidelity DNA Polymerase (New England Biolabs) from the genomic DNA of *E. coli* MG1655, extracted according to Pospiech and Neumann (Pospiech & Neumann, 1995). The PCR fragments were purified using the innuPREP PCR Clean-Up & Gel Extraction Kit (Analytik Jena/iST Innuscreen GmbH, Germany) and cloned into pBAD24 (Guzman et al., 1995) and encoding a C-terminal 6x histidine tag, enabling separation via affinity chromatography. The digestion of vectors and fragments and subsequent ligation (T4 DNA Ligase) were performed according to the New England Biolabs protocol. The constructs were used to transform *E. coli* DH5αλ*pir* cells (Metcalf et al., 1996) for plasmid amplification and maintenance. Plasmid DNA was prepared using the Zyppy Plasmid Miniprep Kit (Zymo Research, USA) according to the manufacturer’s specifications and eluted in 50 μL ddH_2_O. Plasmid concentrations were determined using a spectrophotometer (NanoDrop ND-1000, Peqlab Biotechnologie GmbH, Germany). Plasmids were stored at −20 °C. The constructs and primers used in this study are listed in Tables S2 and S3, respectively.

### 2.3 Generation of deletion mutants

*E. coli* single mutant strains from the Keio collection of mutants were employed (Baba et al., 2006). Because the Δ*gabT* strain was unavailable, it was generated in this study from *E. coli* BW25113 via homologous recombination according to the method used to generate the Keio collection mutants. A double mutant lacking *gabT* and *patA* was generated by removing the cassette as described. The plasmids pRed-ET, 709-FLPe (Quick & Easy *E. coli* Gene Deletion Kit, Gene Bridges GmbH, Germany) and pKD13 (Datsenko & Wanner, 2000) as well as the primers used for the procedure are listed in Tables S2 and S3, respectively.

### 2.4 Sequencing

All plasmid constructs and mutant strains were verified by Sanger sequencing at the Genomics Service Unit (LMU München).

### 2.5 Utilization of AGEs as nitrogen source

*E. coli* BW25113 and the single, double, or triple mutants were inoculated overnight at 37 °C in LB; kanamycin sulfate was added when required at 50 μg/mL. A second overnight culture was prepared by washing the cells in M9 without carbon and nitrogen and diluting them 100-fold in M9 medium with 20 mM D-glucose as the carbon source and 10 mM NH_4_Cl as the nitrogen source. The cells were inoculated to a final OD_600_ of 0.01 in a clear flat-bottom 96-well plate in 200 μL of M9 medium with the following sole nitrogen sources: CML, NH_4_Cl, or no nitrogen. Growth was carried out for 48 hours at 37 °C and a rotational speed of 200 rpm in a plate reader (CLARIOstar Plus, BMG LABTECH, Germany). OD_600_ was measured every 30 minutes. The maximum OD_600_ increase between 0 and 48 hours was reported as the mean of three independent biological replicates, corrected by a factor of 1.6 based on the volume in the wells. Paired t-tests were performed with GraphPad Prism 9.

### 2.6 Protein overexpression and purification

*E. coli* LMG194 cells transformed with pBAD24 encoding a C-terminal histidine-tagged transaminase were grown in LB with 100 µg/mL carbenicillin disodium at 37 °C until an OD_600_ of 0.5 was reached. Overexpression was induced with 0.2% (w/v) L-arabinose and carried out overnight at 18 °C. The cells were harvested by centrifugation (4 °C, 6,240 × *g*, 30 min) and resuspended in purification buffer composed of 20 mM potassium phosphate, 500 mM KCl, 0.1 mM pyridoxal 5’-phosphate (PLP), 1 mM dithiothreitol (DTT), and 20 mM imidazole, modified from previously published methods (Knorr et al., 2018; Liu et al., 2005). The cells were disrupted using ultrasound treatment for 3 cycles of 1.5 min in pulse mode (0.5 s on/off) at an amplitude of 35%. The proteins were purified from the cytosol via Ni^2+^-affinity chromatography (Ni-NTA Agarose, Qiagen). Two wash steps were performed: 20 column volumes (CV) with 20 mM imidazole and 20 CV with 50 mM imidazole, followed by elution with 250 mM imidazole. The chosen fractions were further subjected to size exclusion chromatography using the ÄKTA FPLC system equipped with a Superdex 200 10/300 GL column. The column was equilibrated with 3 CV of water and 2 CV of elution buffer (20 mM potassium phosphate, 500 mM KCl, 0.1 mM PLP, 1 mM DTT, 10% glycerol). The samples were eluted with 1.5 CV, and 10 fractions of 0.5 mL volume were collected. Protein purity was assessed via SDS-PAGE, and concentration was calculated from the A_280_ measured with a spectrophotometer (NanoDrop ND-1000, Peqlab Biotechnologie GmbH, Germany).

### 2.7 Enzymatic assays

The purified transaminases were assayed *in vitro* to determine their specific activity in a luminescence assay based on the production of glutamate. The substrates chosen for this study were CML, CM-Cad (synthesized according to the literature (Hellwig et al., 2019; Hellwig et al., 2011), see Table S4)), α-ketoglutarate, γ-aminobutyric acid (GABA), *N*_α_-acetylornithine (Ac-Orn), L-aspartate, L-isoleucine, L-phenylalanine, and putrescine. The Glutamate-Glo™ Assay kit from Promega was employed. The transaminase reaction mix was composed as follows: 100 mM potassium phosphate, 100 mM KCl, 0.1 mM PLP, 1 mM EDTA, 5 mM α-ketoglutarate, 5 mM substrate (CML, CM-Cad, or cognate substrate), and 0.01 mg/mL transaminase at pH 8.0, modified from (Knorr et al., 2018; Liu et al., 2005). This reaction mix was combined with the Glutamate-Glo reaction mix in a 1:1 ratio. The kit reaction mix contains glutamate dehydrogenase, NAD⁺, reductase, reductase substrate, and luciferin detection solution. The reaction was started by adding the transaminases and followed continuously by measuring luminescence every 5 minutes for up to 15 h in a plate reader (CLARIOstar Plus, BMG LABTECH, Germany). A standard curve including glutamate in the reaction buffer was generated according to the kit protocol. The luminescence over time was plotted, and the linear range was used to calculate the specific activity, defined as μmol_product × min⁻¹ × mg_enzyme⁻¹. Controls including substrate but no enzyme, or enzyme alone in the presence of α-ketoglutarate, were included. The degradation of the reagents and formation of the products were further assessed via HPLC-MS in an *in vitro* time series. The buffer, enzyme, and substrate concentrations were the same as reported above. The reaction was performed in 1.5 mL tubes in a total volume of 1 mL at 37 °C for 1 h for the native substrates and 24 h for CML and CM-Cad. The reaction was started by adding 0.01 mg/mL of enzyme. Samples were collected at defined time points, and the reaction was stopped with an equal volume of 200 mM HCl. The quantification of the metabolites was performed via HPLC-MS. The results are shown as the mean and standard deviation of at least three independent replicates.

### 2.8 Quantification and identification of substrates and products via HPLC-MS/MS and UHPLC-TOF-MS

For quantification of the substrates by tandem mass spectrometry, 5 μL of sample was diluted with 995 μL of aqueous 10 mM nonafluoropentanoic acid (NFPA, Sigma-Aldrich, Germany) solution, centrifuged (10,000 x g, 10 min, room temperature) and transferred to a vial. The analysis was performed on an HPLC-MS/MS system consisting of a binary pump (G7104C), an autosampler (G7129C), a column thermostat (G7116A) and a triple-quadrupole mass spectrometer (G6465BA; all from Agilent Technologies). A stainless-steel column (Zorbax Eclipse Plus C18, 2.1 mm × 100 mm, 3.5 μm, Agilent) was used for separation at a temperature of 35 °C. The mobile phases used were 10 mM NFPA in water as eluent A and 10 mM NFPA in acetonitrile as eluent B, in a gradient (0 min, 2% B; 5 min, 23% B; 6 min, 66% B; 9 min, 66% B; 10 min, 2% B; 14 min, 2% B) at a flow rate of 0.25 mL/min and an injection volume of 10 μL. The mass spectrometer operated in positive MRM mode (dry gas flow, 13 L nitrogen/min; gas temperature, 300 °C; nebulizer pressure, 35 psi; capillary voltage, 4000 V). The parameters for the MRM measurements are shown in Table S5. The same method was used to record product ion scans with varying collision energies (10-30 eV) at preset *m/z* ratios.

To identify the products by TOF-MS, in accordance with a method for the derivatization of α-keto acids (Sajapin & Hellwig, 2020), 50 μL of sample was mixed with 50 μL of a 0.1% (w/v) solution of 2,4-dinitrophenylhydrazine (DNPH, Arcos Organics, Belgium) in 2 M HCl and incubated for 5 minutes at room temperature. This was followed by the addition of 100 μL of 1 M NaOH, 300 μL of acetonitrile and 500 μL of 0.1% aqueous formic acid (Carl Roth, Germany), membrane filtration (0.2 μm) and transfer to a vial. Analysis was performed on an Infinity 1290 UHPLC system (Agilent) consisting of a binary pump with an integrated degasser, an autosampler (G4226A), and a UV detector (G4212A) connected to a TIMS-TOF mass spectrometer (Bruker). The mobile phases used were 0.1% formic acid in water as eluent A and 0.1% formic acid in a mixture of acetonitrile and water (90/10, v/v) as eluent B. Separations with an injection volume of 1 μL were performed at 40 °C on a stainless-steel column filled with the material Eurospher-II 100 (50 mm × 2.0 mm, 3 μm, Knauer) at a flow rate of 0.2 mL/min and using a gradient (0 min, 20% B; 17 min, 67% B; 25 min, 100% B; 30 min, 100% B; 31 min, 20% B; 37 min, 20% B). The mass spectrometer operated in positive scan mode (*m/z* 20-1300; scan time, 500 ms; dry gas flow, 10 L nitrogen/min; dry temperature, 220 °C; nebulizer pressure, 2.2 bar; capillary voltage, 4500 V).

### 2.9 Statistical analysis

Data displayed in this study are shown as means ± standard deviation (SD) of three independent biological replicates, unless otherwise stated. Statistical tests were performed using GraphPad Prism 10 or Excel. To perform a t-test, the unpaired t-test option was chosen. Statistical significance was defined as p < 0.05 (*p < 0.05, **p < 0.01, ***p < 0.001, ****p < 0.0001). Detailed information regarding the statistical test applied and the number of replicates (n) are given in the legends.

## 3. Results

In our previous study, we identified the ornithine decarboxylase SpeC as the enzyme responsible for the initial step in the metabolism of CML, namely its decarboxylation to CM-Cad (Aveta *et al*., 2026). However, comprehensive metabolomic analyses demonstrated the existence of several other carboxymethylated products, pointing toward a more extensive degradation network (Bui et al., 2019; Hellwig et al., 2019). The physiological fact that *E. coli* can utilize CML as a sole nitrogen source confirms that these metabolites are functional intermediates of a catabolic sequence (Aveta et al., 2026). Since the initial SpeC-mediated decarboxylation does not liberate nitrogen, we sought to identify the subsequent enzymatic steps required to channel nitrogen into central metabolism. In view of the slow metabolic turnover of CML and following the logic of our SpeC identification, we hypothesized that the subsequent steps also rely on the promiscuous activity of specialized enzymes (Fig. 1). We focused our search on glutamate-dependent transaminases, which typically serve as central hubs for nitrogen distribution by transferring amino groups to α-ketoglutarate, yielding L-glutamate – the primary nitrogen carrier in the cell (Reitzer, 2003; Walker & van der Donk, 2016).

**Figure 1.**
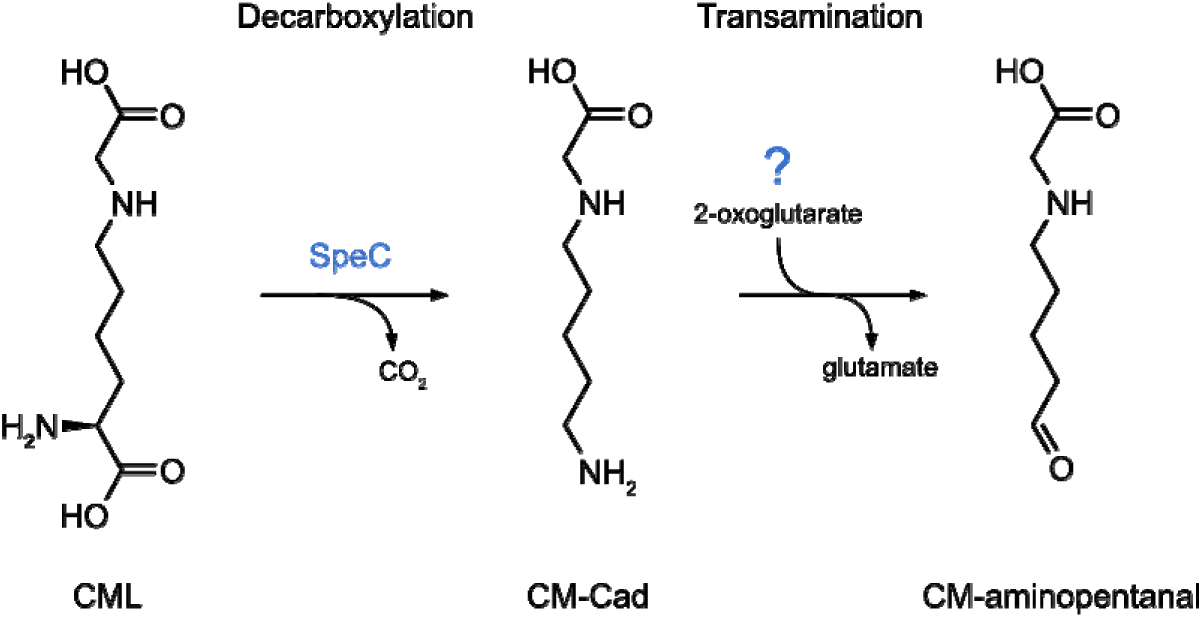
**Proposed metabolism of CML in *E. coli*.**

From the 20 annotated transaminases in the *E. coli* K12 genome (Table S6), we selected eight candidates for characterization, drawing on structural similarities between CML or CM-Cad and their native substrates: ArgD, AspC, AstC, GabT, IlvE, PatA, PuuE, and TyrB (Table 1).

**Table 1.** The transaminase genes of *E. coli* MG1655 selected for this study. The information was collected from the Ecocyc database (Karp Peter et al., 2025)

| Gene | Protein annotation | Substrate |
| --- | --- | --- |
| *argD* | N-acetylornithine aminotransferase/N-succinyldiaminopimelate aminotransferase | N-succinyl-(2S,6S)-2,6-diaminoheptanedioate, N$_{\alpha}$-acetyl-L-ornithine |
| *aspC* | L-aspartate aminotransferase | L-aspartate |
| *astC* | Succinylornithine/acetylornithine transaminase | N-succinylornithine/ N$_{\alpha}$-acetylornithine |
| *gabT* | 4-aminobutyrate aminotransferase | 4-aminobutanoate, 5-aminopentanoate |
| *ilvE* | Branched chain amino acid aminotransferase | L-leucine, L-isoleucine, L-valine, L-phenylalanine |
| *patA* | Putrescine aminotransferase | Putrescine, alkane-$\alpha,\omega$-diamine |
| *puuE* | 4-aminobutyrate aminotransferase | 4-aminobutanoate |
| *tyrB* | Aromatic amino acid aminotransferase | L-tyrosine, L-phenylalanine, L-leucine |

By employing the successful strategy used for the identification of SpeC (Aveta et al., 2026), we evaluated single-gene deletion mutants from the Keio collection (Baba et al., 2006) and assessed growth in M9 minimal medium with CML as the sole nitrogen source. In contrast to the clear phenotype of a *speC* deletion, none of the individual transaminase mutants exhibited a significant growth defect (Fig. S1), nor could we detect any significant changes in the metabolic profile (data not shown).

This result suggested that we either selected the wrong candidate genes or, more likely, highlight a high level of functional redundancy within the *E. coli* transaminase network, where overlapping underground activities mask the contribution of any single enzyme in a cellular context. Building on the latter assumption, we transitioned to a systematic *in vitro* characterization to resolve enzymatic capabilities isolated from a cellular context. We expressed the eight transaminases with a C-terminal His-tag and purified them to homogeneity using Ni-NTA affinity chromatography followed by size-exclusion chromatography (SEC). The purity and identity of the recombinant enzymes were verified by SDS-PAGE (Figs. S2-S3), providing a robust experimental platform for precise enzymatic assays to identify their specific activities (Table S7). Crucially, although the Δ*speC* mutant in our previous study exhibited significantly impaired nitrogen channeling, its growth on CML as a sole nitrogen source was not completely abolished. This pivotal observation suggested that nitrogen liberation does not strictly depend on prior decarboxylation, but might also occur via the direct transamination of CML itself. This may as well explain why the degradation of CML that was observed for *E. coli* wild-type and deletion mutants (Hellwig et al., 2019) could not completely be assigned to the decarboxylation product CM-Cad.

We established an *in vitro* assay utilizing α-ketoglutarate as the primary amino group acceptor and pyridoxal 5′-phosphate (PLP) as the obligate cofactor for our selected transaminases (Bender, 1985). As the successful transamination of the substrates directly yields L-glutamate, we employed the bioluminescent Glutamate-Glo™ assay (Promega) as a highly sensitive screening method to detect active enzyme candidates (Fig. S4). The transaminases were assayed in the presence of CML, CM-Cad, and their respective native substrates (Tab. 1) to establish a baseline of specific activity (Fig. 2A-C, S5-S6). The luminescence output, which is proportional to the amount of L-glutamate formed, revealed that three specific enzymes possess underground activity towards the non-canonical carboxymethylated substrates. IlvE and PatA demonstrated the ability to accept CML as a substrate, exhibiting specific activities of 0.009 and 0.015 μmol min⁻ ¹ mg⁻ ¹ of enzyme, respectively. When evaluating CM-Cad, both PatA and GabT showed transaminase activity, with specific activities of 0.014 and 0.008 μmol min⁻ ¹ mg⁻¹, respectively. Notably, PatA emerged as the most versatile catalyst in this catabolic node, being the only transaminase capable of processing both CML and the decarboxylated derivative CM-Cad. While all enzymes exhibited their expected robust activity toward native substrates (ranging from 0.4 to 0.6 μmol min⁻¹ mg⁻¹), the remaining candidate enzymes (ArgD, AstC, and PuuE) showed only negligible activity toward CML or CM-Cad, with glutamate produced at the threshold of detection.

**Figure 2.**
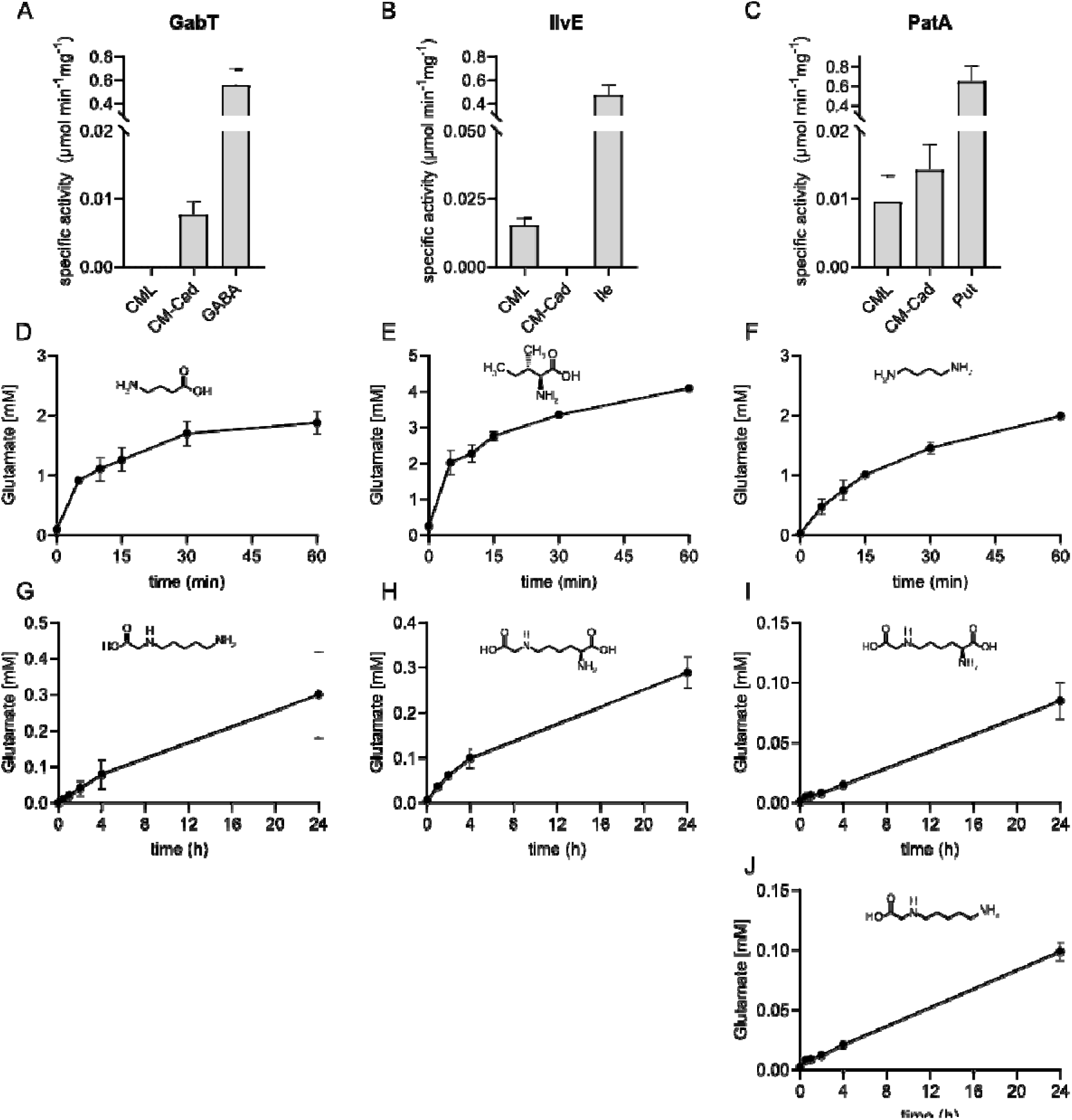
*In vitro* catalytic characterization of the transaminases GabT, IlvE, and PatA. (A–C) Specific transaminase activity of GabT (A), IlvE (B), and PatA (C) determined via the bioluminescent Glutamate-Glo™ assay. The purified enzymes were assayed in the presence of their respective native control substrates γ-aminobutyric acid [GABA], L-isoleucine [Ile], or putrescine [Put]) alongside the non-canonical substrates CML and CM-Cad. Specific activity is expressed as µmol min⁻¹ mg⁻¹ of enzyme. **(D–J)** Time-course quantification of L-glutamate formation determined by HPLC-UV, serving as a direct readout for substrate transamination. The chemical structures of the utilized substrates are depicted within the respective panels. All data points and bar graphs represent the mean ± standard deviation (SD) of three independent replicates (n = 3).

To validate the initial screening results and monitor the stoichiometric conversion over time, we performed continuous time-course measurements using high-performance liquid chromatography coupled with mass spectrometry to quantify L-glutamate formation. For the native control substrates, rapid transamination kinetics were observed within the first 60 minutes of incubation (Fig. 2D-F). By contrast, the conversion of the non-canonical carboxymethylated substrates proceeded at a significantly slower rate, reflecting the character of enzyme promiscuity. Over an extended 24-hour period, continuous HPLC-MS/MS monitoring revealed a steady increase in glutamate levels; GabT yielded approximately 0.3 mM glutamate from 5 mM CM-Cad (Fig. 2G), while IlvE achieved an analogous turnover when provided with 5 mM CML (Fig. 2H). PatA demonstrated its dual specificity by facilitating a continuous conversion of approximately 0.1 mM of both 5 mM CML and CM-Cad to glutamate over the same time frame (Fig. 2I-J). While tracking L-glutamate formation confirmed the continuous transfer of the amino group (Fig. S7), definitive proof of the catabolic pathway required the structural identification of the corresponding signature metabolites derived from the transaminated carbon skeletons (Fig. 3). Therefore, the 24-hour end-point reaction mixtures were subjected to HPLC coupled with tandem mass spectrometry (HPLC-MS/MS). Because the enzymatic transamination of primary amines generates highly reactive and unstable carbonyl compounds – specifically CM-AOHA from CML and CM-APL from CM-Cad – we used 2,4-dinitrophenylhydrazine (DNPH) to trap these transient intermediates as stable hydrazones. MS/MS analysis of the DNPH-derivatized samples directly confirmed the presence of the protonated ion corresponding to the dinitrophenylhydrazone of the linear intermediate CM-AOHA in assays containing PatA and IlvE (Figs. 3 and S8). Interestingly, in underivatized samples from the CML incubations, we identified an additional cyclic product: CM-tetrahydropicolinic acid (CM-THPA), which arises from the spontaneous, non-enzymatic dehydration and intramolecular ring closure of the linear intermediate CM-AOHA (Figs. S9-S10). Similarly, MS/MS analysis of the CM-Cad transamination assays with GabT and PatA detected CM-Pip, resulting from the rapid, spontaneous cyclization of the highly reactive CM-APL intermediate. The direct detection of these signature metabolites, in alignment with the continuous glutamate formation kinetics, provides unambiguous structural and functional proof for the divergent transamination of CML and CM-Cad by IlvE, GabT, and PatA.

**Figure 3.**
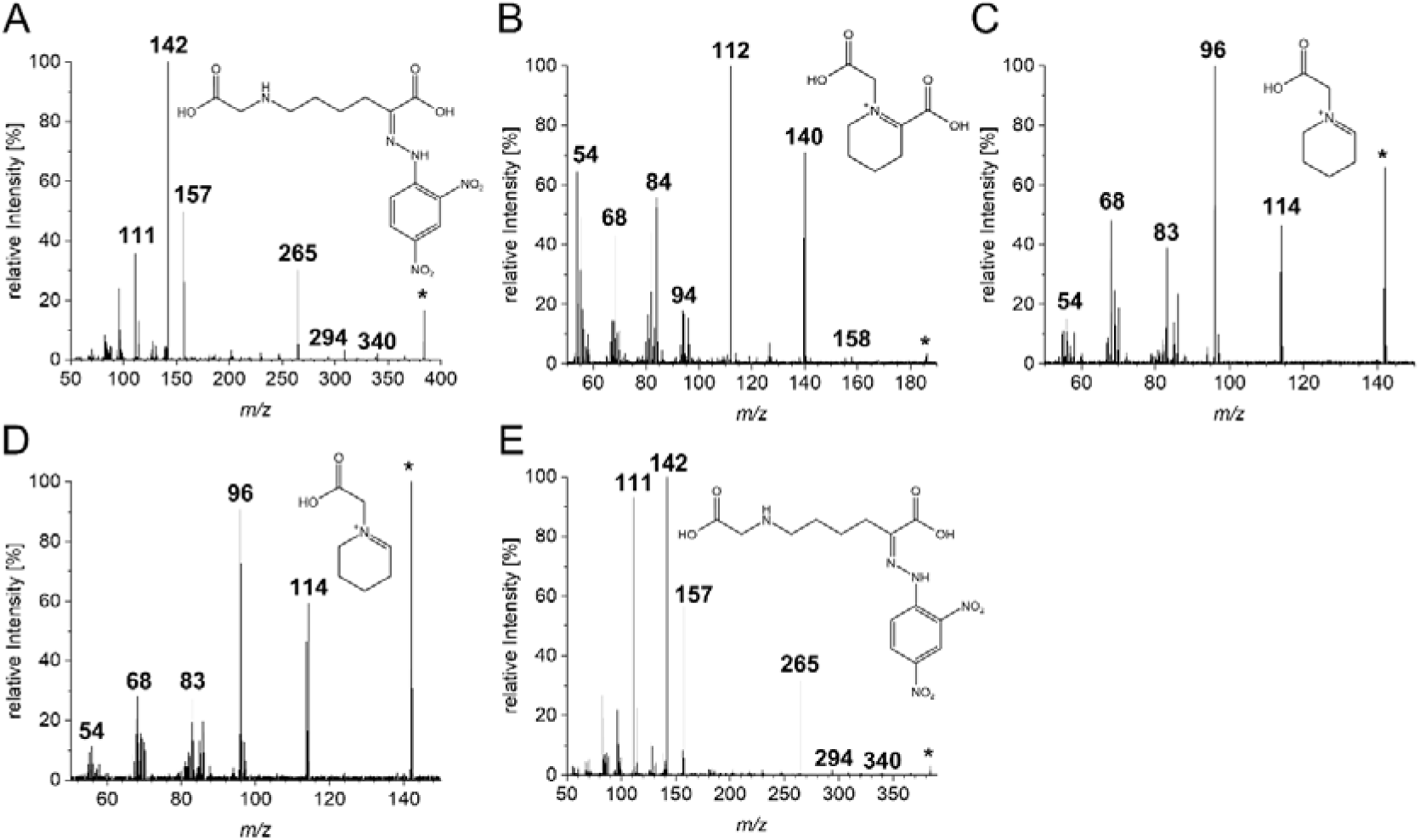
Structural identification of signature transamination products via HPLC-MS/MS. (A, **E)** MS/MS spectra of the DNPH-derivatized linear intermediate CM-AOHA resulting from the transamination of CML by PatA (A) and IlvE (E). **(B)** MS/MS spectrum of the underivatized, spontaneously cyclized product CM-THPA resulting from the incubation of CML with PatA. **(C, D)** MS/MS spectra of the spontaneously formed CM-Pip resulting from the transamination of CM-Cad by PatA (C) and GabT (D). The proposed chemical structures of the parent ions are depicted within the respective panels.

The *in vitro* catalytic profiles validated the initial hypothesis that a redundant transaminase network, characterized by overlapping underground activities, masks the contributions of individual enzymes *in vivo*. Consequently, we postulated that only the simultaneous inactivation of these specific candidates would collapse the network redundancy and expose the metabolic block within the cellular context. This rationale guided the construction of combinatorial deletion mutants (Fig. 4).

**Figure 4.**
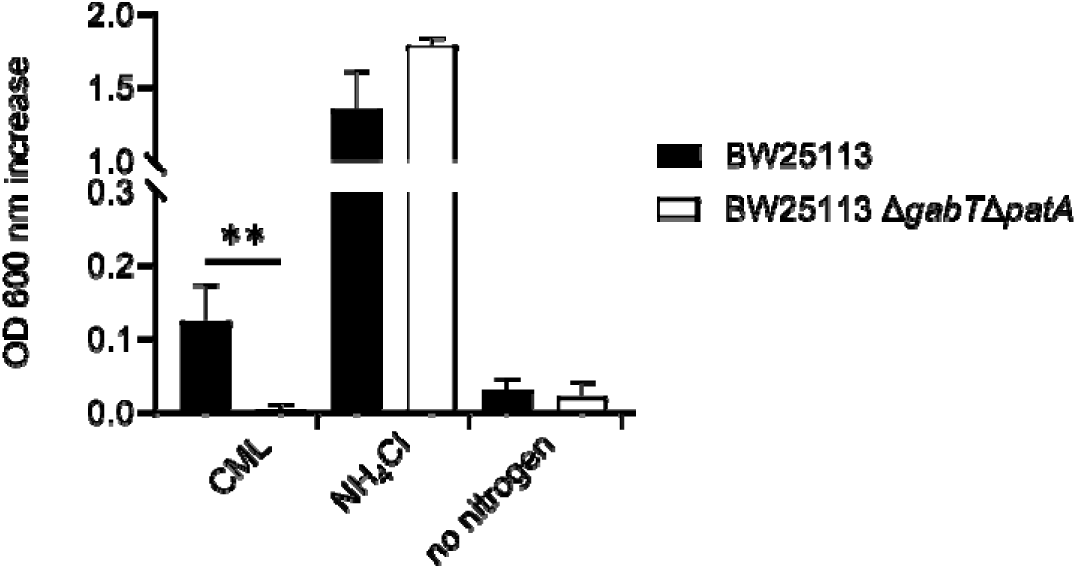
*In vivo* effect of the double transaminase mutants. Growth analysis of *E. coli* BW25113 and the double mutant Δ*gabT*Δ*patA* in M9 minimal medium with 10 mM CML, NH_4_Cl or without a nitrogen source. Depicted is the maximum increase in OD_600_ over 48 h of incubation at 37 °C. The data are shown as mean and SD of three independent replicates (n = 3), and Unpaired *t*-test was performed with GraphPad Prism 10, yielding a *p-*value of 0.0060.

However, the inclusion of *ilvE* in this *in vivo* strategy is technically precluded. The *ilvE* gene encodes the primary branched-chain amino acid aminotransferase, and its deletion intrinsically causes an auxotrophy for isoleucine, leucine, and valine (Amorim Franco & Blanchard, 2017). Cultivating an *ilvE* mutant necessitates their supplementation to the minimal medium. These amino acids would then also serve as alternative nitrogen sources and their addition would critically confound the growth assay, which strictly relies on CML as the sole source of nitrogen. Thus, we focused the *in vivo* phenotypic evaluation exclusively on the Δ*gabT* Δ*patA* double mutant. Phenotypic quantification of this mutant in minimal medium supplemented with CML as the sole nitrogen source confirmed the hypothesis: While the wild-type strain exhibited robust growth, the simultaneous disruption of the two transaminase genes completely abolished bacterial growth (Fig. 4). This absolute defect stands in sharp contrast to the previously characterized Δ*speC* mutant. While the elimination of the decarboxylase SpeC severely bottlenecked nitrogen channeling, it did not entirely eradicate growth, leaving a residual capacity for CML utilization.

Conversely, the complete growth arrest of the transaminase double mutant demonstrates that no alternative metabolic bypass or secondary underground pathway is capable to compensate for the lack of these two enzymes. To rule out generalized metabolic deficiencies or fitness costs introduced by the multi-gene deletions, growth was monitored using ammonium chloride as a readily assimilable positive control. Here, the growth kinetics and final biomass yield of the double mutant were completely indistinguishable from the parental wild-type strain. Together, these *in vivo* findings demonstrate that the concerted action of GabT and PatA represents a critical functional node for CML-derived nitrogen utilization. While the potential physiological contribution of IlvE remains to be fully defined due to growth-supplementation constraints, the absolute growth arrest of the double mutant highlights that GabT and PatA are indispensable components within this interconnected transamination network under the tested conditions.

## 4. Concluding discussion

The present study elucidates the central enzymatic steps of nitrogen assimilation from the AGE CML in *E. coli*. Building upon the identification of the ornithine decarboxylase SpeC as the catalyst for the initial degradation to CM-Cad, our data define PatA, GabT, and IlvE as the responsible transaminases for the subsequent transamination. These reactions transfer the nitrogen to α-ketoglutarate, feeding it into central metabolism as L-glutamate, while the highly reactive carbon skeletons spontaneously condense into the cyclic compounds CM-THPA and CM-Pip.

This opportunistic utilization of the AGE CML expands the paradigm of bacterial underground metabolism and stands in stark contrast to the catabolism of early glycation products (Graf von Armansperg et al., 2021; Lassak et al., 2023). While the mammalian host efficiently absorbs up to 95% of canonical dietary protein in the upper gastrointestinal tract, the unabsorbed fraction reaching the distal gut is disproportionately enriched with modified amino acids – including both early Maillard reaction intermediates and AGEs. However, irrespective of this shared mechanism of luminal enrichment, the absolute dietary intake of CML is significantly lower than that of early glycation products (Hellwig et al., 2024; Snelson & Coughlan, 2019). Driven by the substantial intake and high nutritional value of accumulating early glycation products like fructoselysine, bacteria have evolved highly specific, dedicated genetic loci, such as the *frl* operon, for their efficient metabolism (Graf von Armansperg et al., 2021; Lassak et al., 2023). By contrast, the comparatively lower abundance of CML does not induce specialized catabolic operons. Instead, *E. coli* prevents these unabsorbed diet constituents from being excreted by relying on the promiscuous repertoire of its existing enzymes. Regardless of their divergent specificities for canonical substrates, PatA, GabT, and IlvE exhibit a converging promiscuity that allows them to process these sterically demanding, carboxymethylated derivatives. This functional redundancy represents a robust evolutionary mechanism to overcome nutrient limitations by exploiting atypical nitrogen sources without requiring time-consuming genomic adaptations.

The physiological relevance of this network was validated *in vivo* through the combinatorial inactivation of GabT and PatA, which led to a complete growth arrest on CML. This result demonstrates that despite the broad catalytic capabilities of the *E. coli* transaminome, specific functional nodes exist whose disruption completely blocks the catabolic flux. While the absolute *in vivo* contribution of IlvE remains partially obscured due to inherent auxotrophy and supplementation constraints, the essentiality of the GabT/PatA node is unambiguously established under the tested conditions.

While the initial steps of nitrogen liberation are now resolved, the complete structural fate of the deaminated carbon skeletons requires further investigation. Previous *in vivo* metabolic profiling has identified CM-APA as a prominent CML degradation product (Hellwig et al., 2019). Although our current systematic metabolomic setup did not specifically show CM-APA formation, it is highly probable that the transient, highly reactive aldehyde intermediates generated by transamination do not exclusively undergo spontaneous cyclization. The oxidation of these intermediates by an as-yet-unidentified dehydrogenase to yield CM-APA represents a logical subsequent and presumably last enzymatic step. Identifying the specific dehydrogenases responsible for this conversion will be essential to fully map the downstream branches of the CML degradation network.

Furthermore, the physiological fate of these terminal metabolites is of significant clinical interest. CML is implicated in age related pathologies (Hellwig et al., 2024; Mossad et al., 2022). Future studies must evaluate whether microbial transformation products like CM-Pip or CM-APA possess any toxicological relevance for the human host, or conversely, whether this bacterial degradation pathway provides a protective clearance mechanism that limits the intestinal transit and potential residual absorption of exogenous AGEs.

In conclusion, this work provides mechanistic proof for a branched, promiscuity-driven transaminase network. It demonstrates at the molecular level how members of the microbiome leverage their existing biochemical architecture to process anthropogenic molecules. The identification of PatA and GabT as essential nodes of CML catabolism provides the foundation for future targeted analyses of microbial AGE metabolism and its systemic effects on the host-microbiome axis.

## Acknowledgements

None.

## 5. Statements and declarations

### Author Contributions

The manuscript was written by JL, MH, PV and EFA. The study was designed by JL and MH, with contribution of all the authors. The *in vivo* experiments were performed by EFA. The *in vitro* biochemical characterization of *E. coli* transaminases was performed by EFA, PV, and VH. The synthesis of the compounds used in this study was performed by PV. The HPLC-MS/MS measurements and analysis were performed by PV, MH and VH. All authors have given approval to the final version of the manuscript.

### Notes

The authors declare no competing financial interest.

## Chemical abbreviations

AGE: advanced glycation end-product
CML: Nε-carboxymethyllysine
CM-Cad: N-carboxymethylcadaverine
CM-APA: N-carboxymethyl-aminopentanoic acid
CM-Pip: N-carboxymethyl piperideinium ion
CM-THPA: N-carboxymethyl-tetrahydropicolinic acid
CM-AOHA: N-carboxymethylamino-oxohexanoic acid
CM-APL: N-carboxymethyl-aminopentanal
CM-APA: N-carboxymethyl-aminopentanoic acid

