## Supplementary material for "Divergent specificity of PatA, GabT, and IlvE defines the branched transamination of *N*_ε_-carboxymethyllysine and its metabolite *N*_ε_-carboxymethylcadaverine in *Escherichia coli*"

**- Supporting information -**

Table S1. Strains used in this study.

| **Strain** | **Genotype or description** | **Purpose** | **Source or reference** |
| --- | --- | --- | --- |
| *E. coli* BW25113 | F⁻ Δ(araD-araB)567 Δ(lacZ)4787(::rrnB-3) λ⁻ rph-1 Δ(rhaD-rhaB)568 hsdR514 | in vivo CML degradation | (Datsenko & Wanner, 2000) |
| *E. coli* JW0911 | BW25113, Δ*aspC*::*kan* | in vivo CML degradation | (Baba et al., 2006) |
| *E. coli* JW1737 | BW25113, Δ*aspC*::*kan* | in vivo CML degradation | (Baba et al., 2006) |
| *E. coli* JW3322 | BW25113, Δ*argD*::*kan* | in vivo CML degradation | (Baba et al., 2006) |
| *E. coli* JW5606 | BW25113, Δ*ilvE*::*kan* | in vivo CML degradation | (Baba et al., 2006) |
| *E. coli* JW5510 | BW25113, Δ*patA*::*kan* | in vivo CML degradation | (Baba et al., 2006) |
| *E. coli* JW1295 | BW25113, Δ*puuE*::*kan* | in vivo CML degradation | (Baba et al., 2006) |
| *E. coli* JW4014 | BW25113, Δ*tyrB*::*kan* | in vivo CML degradation | (Baba et al., 2006) |
| *E. coli* JW0593 | BW25113, Δ*ybdL*::*kan* | in vivo CML degradation | (Baba et al., 2006) |
| *E. coli* BW25113 Δ*gabT* | BW25113, Δ*gabT*::*kan* | in vivo CML degradation | this study |
| *E. coli* BW25113 Δ*gabT*Δ*patA* | BW25113, Δ*gabT*Δ*patA*::*kan* | in vivo CML degradation | this study |
| *E. coli* LMG194 | F- Δ(lacIPOZY)X74 galE galK thi rpsL ΔphoA ara714 | protein purification | (Guzman et al., 1995) |
| *E. coli* DH5α λ*pir* | F⁻ φ80dlacZΔM15 Δ(lacZYA-argF)U169 recA1 endA1 hsdR17(rK⁻, mK⁺) phoA supE44 λ⁻ thi-1 gyrA96 relA1 [λ pir] | cloning | (Metcalf et al., 1996) |

Table S2. Plasmids used in this study.

| **Plasmid** | **Purpose** | **Selective marker** | **Reference** |
| --- | --- | --- | --- |
| pBAD24 | Bacterial expression vector; arabinose inducible araBAD promoter (PBAD) | Ampicillin resistance | (Guzman et al., 1995) |
| pBAD24-*aspC* | Purification of *E. coli* AspC | Ampicillin resistance | this study |
| pBAD24-*argD* | Purification of *E. coli* ArgD | Ampicillin resistance | this study |
| pBAD24-*astC* | Purification of *E. coli* AstC | Ampicillin resistance | this study |
| pBAD24-*gabT* | Purification of *E. coli* GabT | Ampicillin resistance | this study |
| pBAD24-*ilvE* | Purification of *E. coli* IlvE | Ampicillin resistance | this study |
| pBAD24-*patA* | Purification of *E. coli* PatA | Ampicillin resistance | this study |
| pBAD24-*puuE* | Purification of *E. coli* PuuE | Ampicillin resistance | this study |
| pBAD24-*tyrB* | Purification of *E. coli* TyrB | Ampicillin resistance | this study |
| pBAD24-*ybdL* | Purification of *E. coli* YbdL | Ampicillin resistance | this study |
| pKD13 | Template for generation of a kanamycin resistance cassette | Kanamycin resistance | (Datsenko & Wanner, 2000) |
| pRed/ET | Recombinase bearing plasmid for insertion of a kanamycin resistance cassette | Ampicillin resistance | Quick and Easy E. coli Gene Deletion Kit, Gene Bridges, Heidelberg |
| 709-FLPe | FLP-recombinase expression plasmid for helper-free excision of resistance cassettes | Ampicillin resistance | Quick and Easy E. coli Gene Deletion Kit, Gene Bridges, Heidelberg |

Table S3. Primers used in this study. The check primers were used to generate a cassette for double homologous recombination using the Keio collection mutants gDNA as a template for *patA* but not for *gabT*, for which a mutant was not available on the collection. This mutant was generated with deletion primers using pKD13 as template.

| **Primer** | **Sequence (5' 3')** |
| --- | --- |
| pBAD24 insert check forward | GGC GTC ACA CTT TGC TAT GC |
| pBAD24 insert check reverse | CAG TTC CCT ACT CTC GCA TG |
| pBAD24 *aspC* cloning forward XbaI-His6-GS-aspC | CTA GAT TAG TGA TGG TGA TGG TGA TGA CTG CCC AGC ACT GCC ACA ATC GCT T |
| pBAD24 *aspC* cloning reverse PciI-aspC | CAT GTT TGA GAA CAT TAC CGC C |
| pBAD24 *astC* cloning forward XbaI-His6-GS-astC | CTA GAT CAG TGA TGG TGA TGG TGA TGT GAT GAA CCT CGG CTA ACA AAG |
| pBAD24 *astC* cloning reverse NcoI-astC | CAT GGG TTC TCA GCC AAT TAC GCG TGA AAA CTT TG |
| pBAD24 *argD* cloning forward XbaI-argD | ATA ATT TCT AGA CGT ATT TAA TGG TGG TGA TGG TGA TGA CTA CCC GCC CCA ACC ACC T |
| pBAD24 *argD* cloning reverse NcoI-argD | CAT GGC AAT TGA ACA AAC AGC AAT TAC |
| pBAD24 *gabT* cloning forward NcoI-gabT | TGC CCA TGG GCA ACA GCA ATA AAG AGT TAA TGC AGC G |
| pBAD24 *gabT* cloning reverse XbaI-His6-GS-gabT | GGA GTC TAG ACT AAT GGT GAT GGT GAT GGT GAT GGC CAC CAC CCT GCT TCG CCT CAT CAA AAC ACT G |
| pBAD24 *ilvE* cloning forward NcoI-ilvE | ATA AAC CAT GGG TAA CAC GAA GAA AGC TGA TTA C |
| pBAD24 *ilvE* cloning reverse XbaI-His6-GS-ilvE | TGT ATT CTA GAT TAA TGG TGA TGG TGA TGG TGA CTG CCT TGA TTA ACT TGA TCT AAC CAG CC |
| pBAD24 *patA* cloning forward NcoI-patA | GCA TAC CAT GGG TTT GAA CAG GTT ACC TTC GAG C |
| pBAD24 *patA* cloning reverse XbaI-His6-GS-patA | GAT TTC TAG ATT AGT GAT GGT GAT GGT GAT GAC TAC CCG CTT CTT CGA CAC TTA CTC G |
| pBAD24 *puuE* cloning forward PciI-PuuE | TGA AAC ATG TCT AGC AAC AAT GAA TTC CAT CAG |
| pBAD24 *puuE* cloning reverse XbaI-His6-GS-PuuE | GCG TTT CTA GAT TAG TGA TGG TGA TGG TGA TGA CTG CCA TCG CTC AGC GCA TCC TG |
| pBAD24 *tyrB* cloning forward NcoI-tyrB | CCA TCC CAT GGG GTT TCA AAA AGT TGA CGC CTA CGC TGG |
| pBAD24 *tyrB* cloning reverse XbaI-His6-GS-tyrB | TGC ATC TAG ATC AGT GAT GGT GAT GGT GAT GTG ATG ACA TCA CCG CAG CAA ACG CC |
| pBAD24 *ybdL* cloning forward NcoI-ybdL | CAT GGG TAC AAA TAA CCC TCT GAT TCC AC |
| pBAD24 *ybdL* cloning reverse XbaI-His6-GS-tyrB | CTA GAC TAA TGG TGA TGA TGG TGG TGA CTA CCA AGC TGG CGC AGG CGT TCA GC |
| *gabT* deletion forward | CTT AGA AAT CAA ATA TAT GTG CAT CGG TCT TTA ACT GGA GAA TGC GAA TGA TTC CGG GGA TCC GTC GAC C |
| *gabT* deletion reverse | GCA GCG TCG CCT CCG GCA TAG GAG CGG CGC TAC TGC TTC GCC TCA TCA AAT GTA GGC TGG AGC TGC TTC G |
| *patA* deletion check forward | CGC AGC AAT CAT CAA ATC CAT ACC CG |
| *patA* deletion check reverse | CGT GAC CAG CCT GGT GCC GTA C |
| *gabT* deletion check forward | CGC TGG AGT ACG GCA TCG TCG |
| *gabT* deletion check reverse | AGC GGC TCC GCC TTA CCC G |

Table S4. Analytical data for characterization of CML and CM-Cad utilized in this study (Aveta et al., 2026).

| **Analytical data for CML** |  |
| --- | --- |
| ^1^H NMR (300 MHz, D_2_O), δ [ppm]: | 1.45 (m, 2H, H-4); 1.70 (m, 2H, H-5); 1.90 (m, 2H, H-3); 3.04 (t, 2H, H-6, J = 7.8); 3.84 (s, 2H, H-1’) 3.98 (t, 1H, J = 6.3 Hz, H-2). |
| HPLC-MS/MS | t_R_, 7.4 min; fragmentation (80 V, 20 eV) of [M + H]^+^ (*m/z* 205) 84 (100), 130 (6), 142 (5). |
| Elemental analysis | Calcd for C_8_H_16_N_2_O_4_ (MW = 204.22): C, 47.05; H, 7.90; N, 13.72. Found: C, 30.95; H, 6.28; N, 9.10; content 66.3 % based on nitrogen. |
| **Analytical data for CM-Cad** |  |
| ^1^H NMR (300 MHz, D_2_O), δ [ppm]: | 1.39 (m, 2H, H-3); 1.66 (m, 4H, H-2, H-4); 2.94 (t, 2H, J = 6.9 Hz, H-1); 3.05 (t, 2H, J = 7.6 Hz, H-5); 3.83 (s, 2H, H-1‘). |
| HPLC-MS/MS | t_R_, 9.0 min; fragmentation (80 V, 20 eV) of [M + H]^+^ (*m/z* 161) 86 (100), 98 (61), 69 (18). |
| Elemental analysis | Calcd for C_7_H_16_N_2_O_2_ (MW = 160.21): C, 52.48; H, 10.07; N, 17.48. Found: C, 19.93; H, 4.68; N, 6.63; content 37.9 % based on nitrogen. |

Table S5. Transitions recorded during MRM measurement of CML, CM-Cad, glutamate, GABA, isoleucine, putrescine.

| **Time window** | **Analyte** | **Transition** | **Fragmentor Voltage** | **Collision energy** | **Q/q** |
| --- | --- | --- | --- | --- | --- |
| 1.5–2.5 min | glutamate | *m/z* 148 → 56 | 80 V | 25 eV | q |
|  |  | *m/z* 148 → 84 | 80 V | 20 eV | Q |
|  |  | *m/z* 148 → 102 | 80 V | 10 eV | q |
| 2.5–6 min | CML | *m/z* 205 → 84 | 80 V | 20 eV | Q |
|  |  | *m/z* 205 → 130 | 80 V | 10 eV | q |
|  |  | *m/z* 205 → 142 | 80 V | 10 eV | q |
|  | GABA | *m/z* 104 → 45 | 80 V | 20 eV | q |
|  |  | *m/z* 104 → 69 | 80 V | 15 eV | q |
|  |  | *m/z* 104 → 87 | 80 V | 10 eV | Q |
| 6–9 min | CM-Cad | *m/z* 161 → 69 | 80 V | 15 eV | q |
|  |  | *m/z* 161 → 86 | 80 V | 10 eV | Q |
|  |  | *m/z* 161 → 98 | 80 V | 20 eV | q |
|  | isoleucine | *m/z* 132 → 57 | 80 V | 30 eV | q |
|  |  | *m/z* 132 → 69 | 80 V | 20 eV | q |
|  |  | *m/z* 132 → 86 | 80 V | 10 eV | Q |
|  | putrescine | *m/z* 89 → 55 | 80 V | 25 eV | q |
|  |  | *m/z* 89 → 72 | 80 V | 10 eV | Q |

Table S6. Genes annotated as transaminases in *E. coli* MG1655, extracted from NCBI.

| **GeneID** | **Symbol** | **description** |
| --- | --- | --- |
| 948296 | *wecE* | dTDP-4-dehydro-6-deoxy-D-glucose transaminase |
| 946255 | *astC* | succinylornithine transaminase |
| 945211 | *ybdL* | methionine transaminase |
| 948278 | *ilvE* | branched-chain-amino-acid aminotransferase |
| 945553 | *aspC* | aspartate aminotransferase |
| 948067 | *gabT* | 4-aminobutyrate aminotransferase GabT |
| 948241 | *glmS* | L-glutamine--D-fructose-6-phosphate aminotransferase |
| 945376 | *bioA* | adenosylmethionine-8-amino-7-oxononanoate aminotransferase |
| 948087 | *avtA* | valine--pyruvate aminotransferase |
| 948563 | *tyrB* | tyrosine aminotransferase |
| 947864 | *argD* | N-acetylornithine aminotransferase/N-succinyldiaminopimelate aminotransferase |
| 945527 | *serC* | phosphoserine/phosphohydroxythreonine aminotransferase |
| 946551 | *hisC* | histidinol-phosphate aminotransferase |
| 945446 | *puuE* | 4-aminobutyrate aminotransferase PuuE |
| 947390 | *gcvT* | aminomethyltransferase |
| 947120 | *patA* | putrescine aminotransferase |
| 947375 | *arnB* | UDP-4-amino-4-deoxy-L-arabinose aminotransferase |
| 946772 | *alaA* | glutamate--pyruvate aminotransferase AlaA |
| 946850 | *alaC* | glutamate--pyruvate aminotransferase AlaC |
| 947875 | *frlB* | fructoselysine 6-phosphate deglycase |

Table S7. Molecular weights of the His-tagged proteins of this study, computed with protparam (Gasteiger et al., 2005)

| **Protein** | **Molecular weight (kDa) (oligomer)** | **Molecular weight (kDa) (monomer)** |
| --- | --- | --- |
| ArgD | 89.47 | 44.73 |
| AspC | 96.63 | 48.32 |
| AstC | 89.08 | 44.54 |
| GabT | 186.97 | 46.74 |
| IlvE | 210.36 | 35.06 |
| PatA | 101.26 | 50.63 |
| PuuE | 182.78 | 45.70 |
| TyrB | 89.01 | 44.50 |

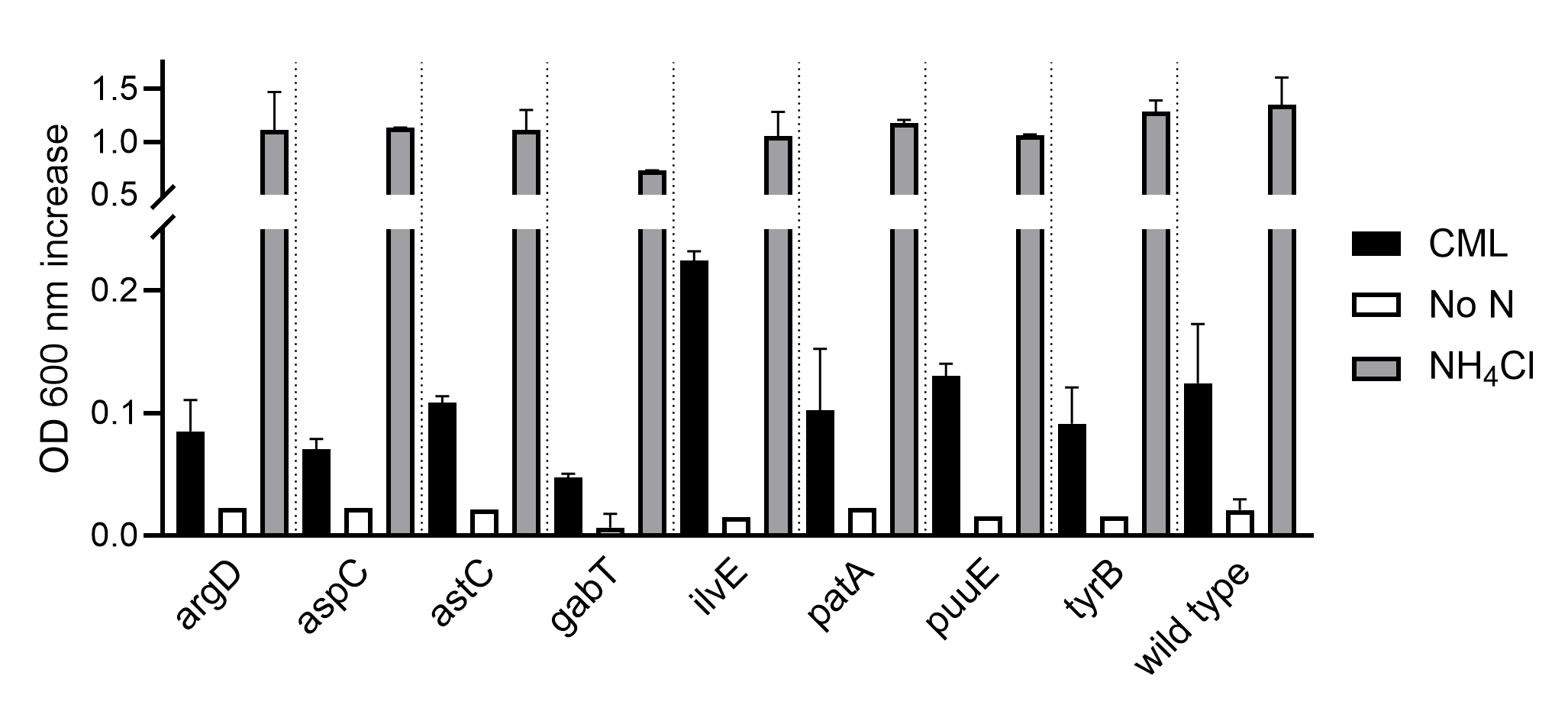

Figure S1. Growth of the *E. coli* BW25113 single transaminase mutants in minimal medium in presence of 10 mM CML, NH_4_Cl or no nitrogen source. The growth is depicted as maximum OD_600_ increase of a 48 h incubation at 37°C. The data are showcased as mean and SD of at least two independent biological replicates (n = 2) except for the wild type (n = 3).

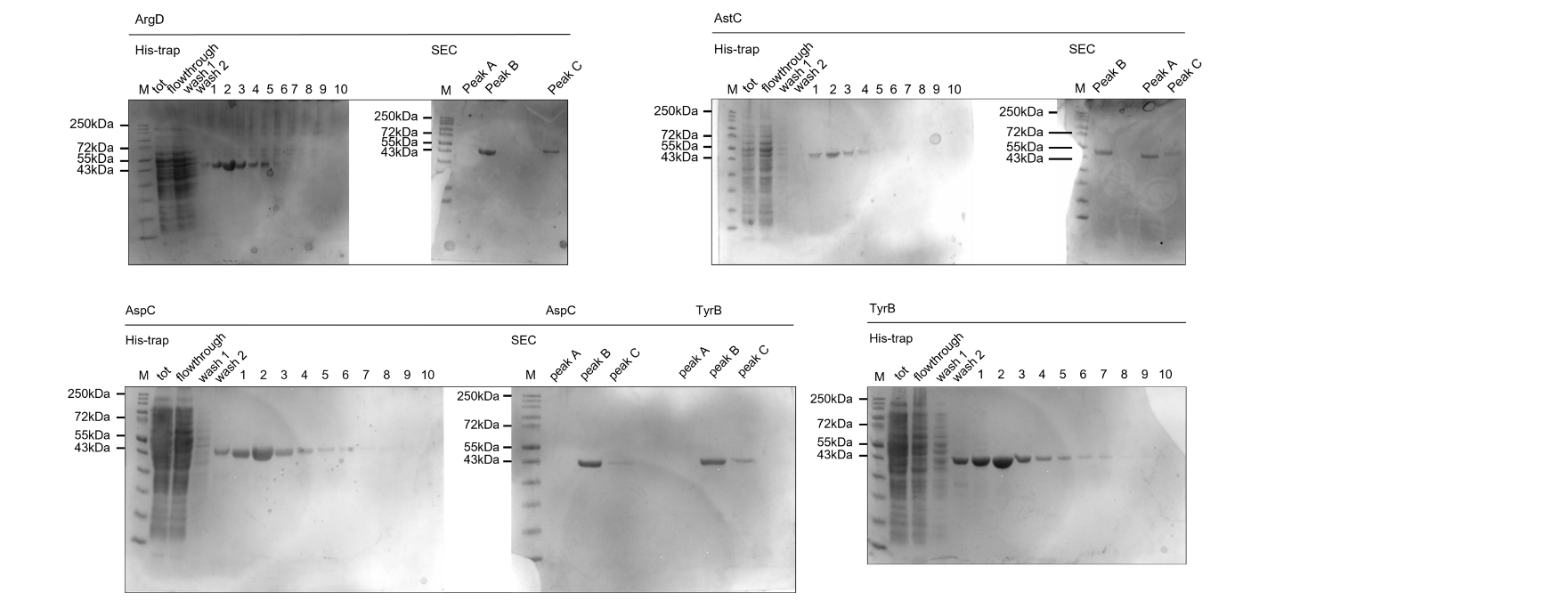

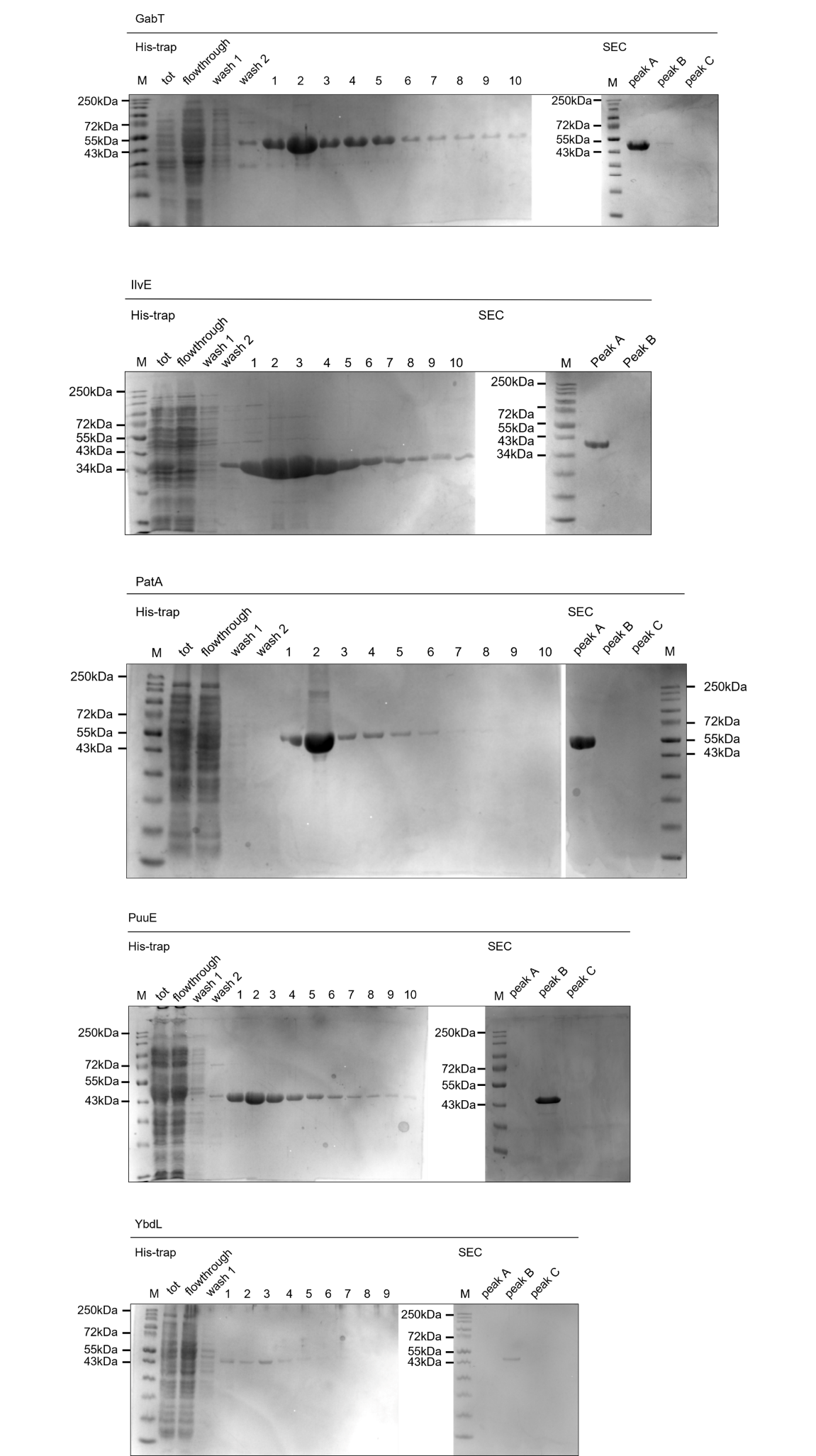

Figure S2. SDS-PAGE of the purified proteins. At various steps of the purification process, 10 μL of sample were collected and combined with Laemmli sample buffer, incubated at 95 °C for 10 minutes and loaded on a 12.5% Tris-glycine SDS-PAGE gel. The proteins were visualized via Instant Coomassie staining (BioRad). Size-exclusion peak fractions correspond to the peaks in figure S3. The molecular weight marker (M) used is the Color Prestained Protein Standard, Broad Range (10-250 kDa, New England Biolabs).

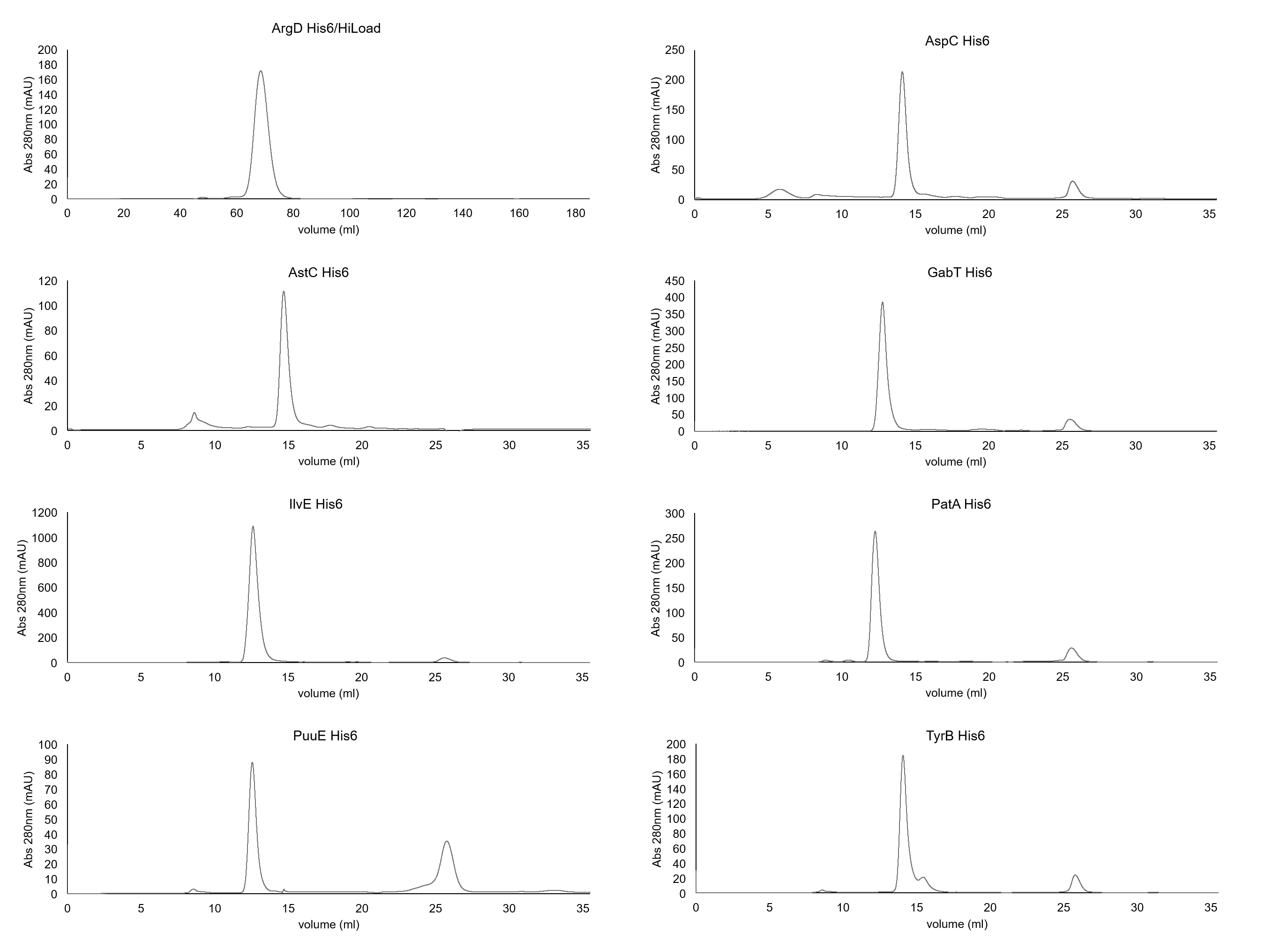

Figure S3. Size exclusion chromatograms of purified transaminases. All the proteins were purified with a Superdex 200 Increase 10/300 GL column, ArgD was purified with the HiLoad version. The fractions containing peaks were checked by SDS-PAGE as shown in figure S2.

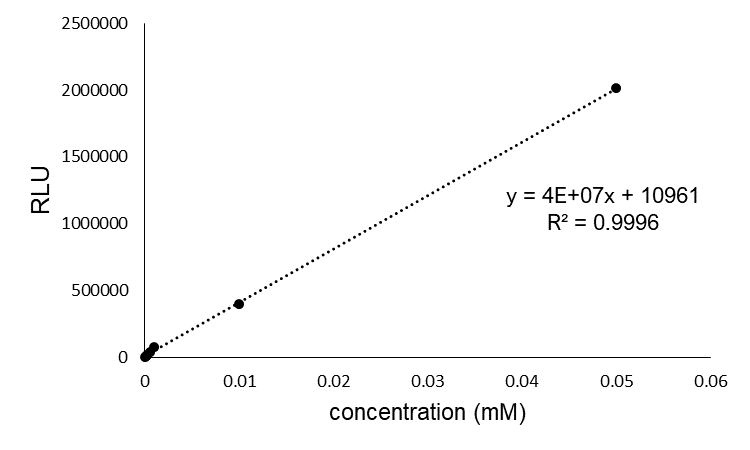

Figure S4. Glutamate-Glo assay standard curve used to determine the specific activity of the transaminases, obtained by plotting Relative Luminescence Units (RLU) against glutamate concentration and performing linear regression using Excel. The standard curve was obtained using the reaction buffer as the glutamate diluent, according to the kit instructions.

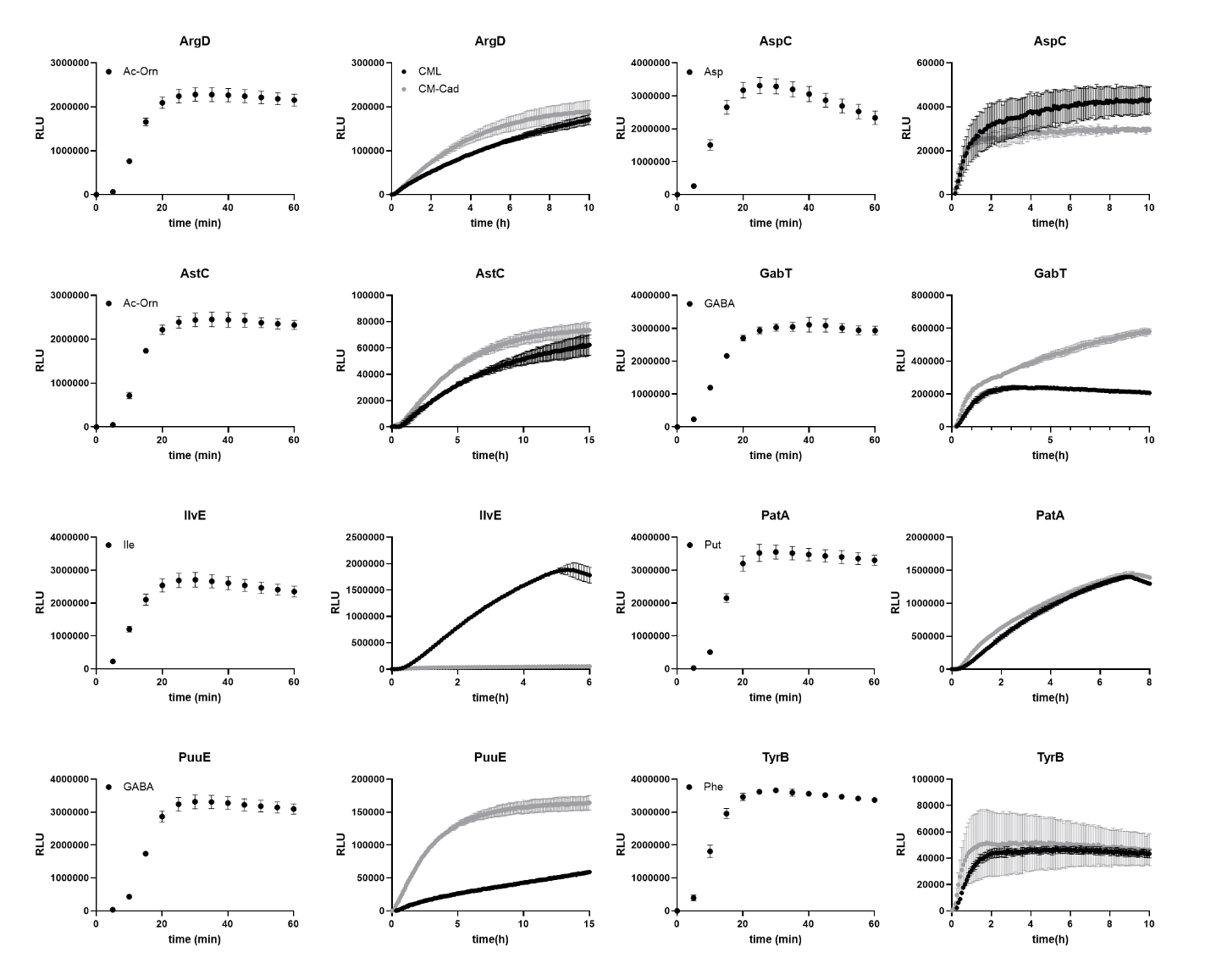

Figure S5. Signal curve obtained from the Glutamate-Glo assay. For each enzyme, the cognate substrate, CML and CM-Cad were included, together with α-ketoglutarate, and the luminescence (RLU) over time was recorded in a plate reader. Abbreviations: *N*_α_-Acetyl-ornithine (Ac-Orn), aspartate (Asp), γ-amino butyrate (GABA), isoleucine (Ile), putrescine (Put), phenylalanine (Phe)The data are shown as mean and SD of three independent replicates (n = 3).

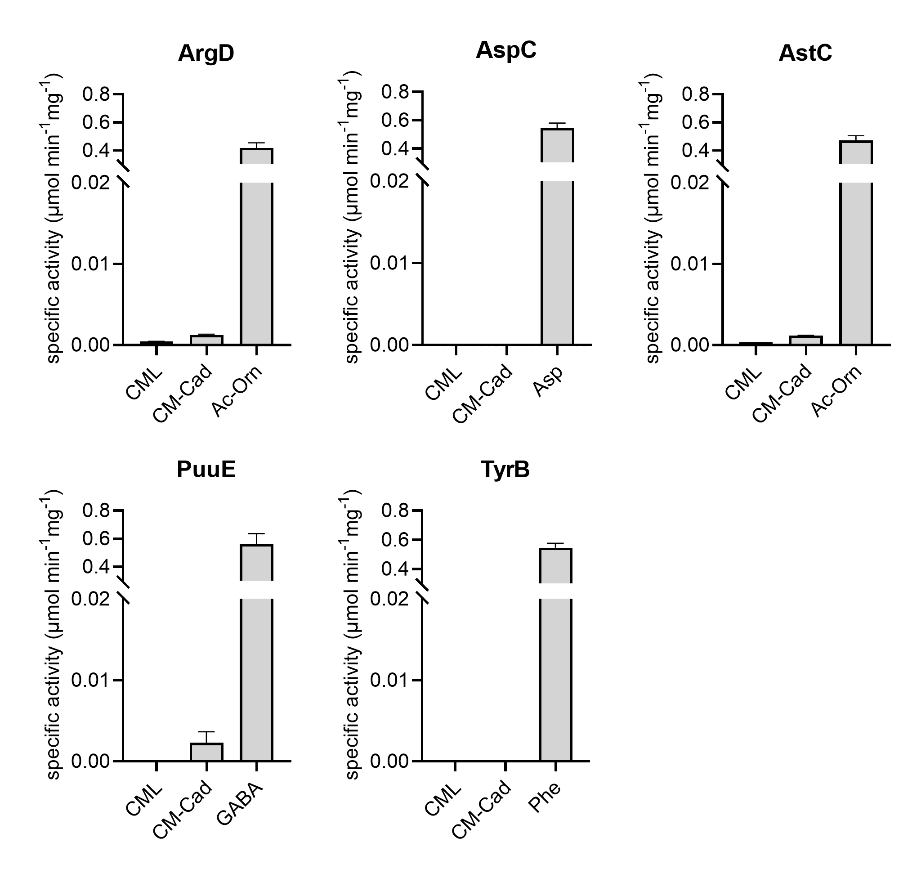

Figure S6. Specific activity of the transaminases determined for CML, CM-Cad and one respective native substrate, calculated from the signal curves in figure S5. The specific activity is expressed as µmol*min^-1^*mg^-1^_enzyme_. Abbreviations: *N*_α_-Acetyl-ornithine (Ac-Orn), aspartate (Asp), γ-amino butyrate (GABA), phenylalanine (Phe) The data are shown as mean and SD of three independent replicates (n = 3).

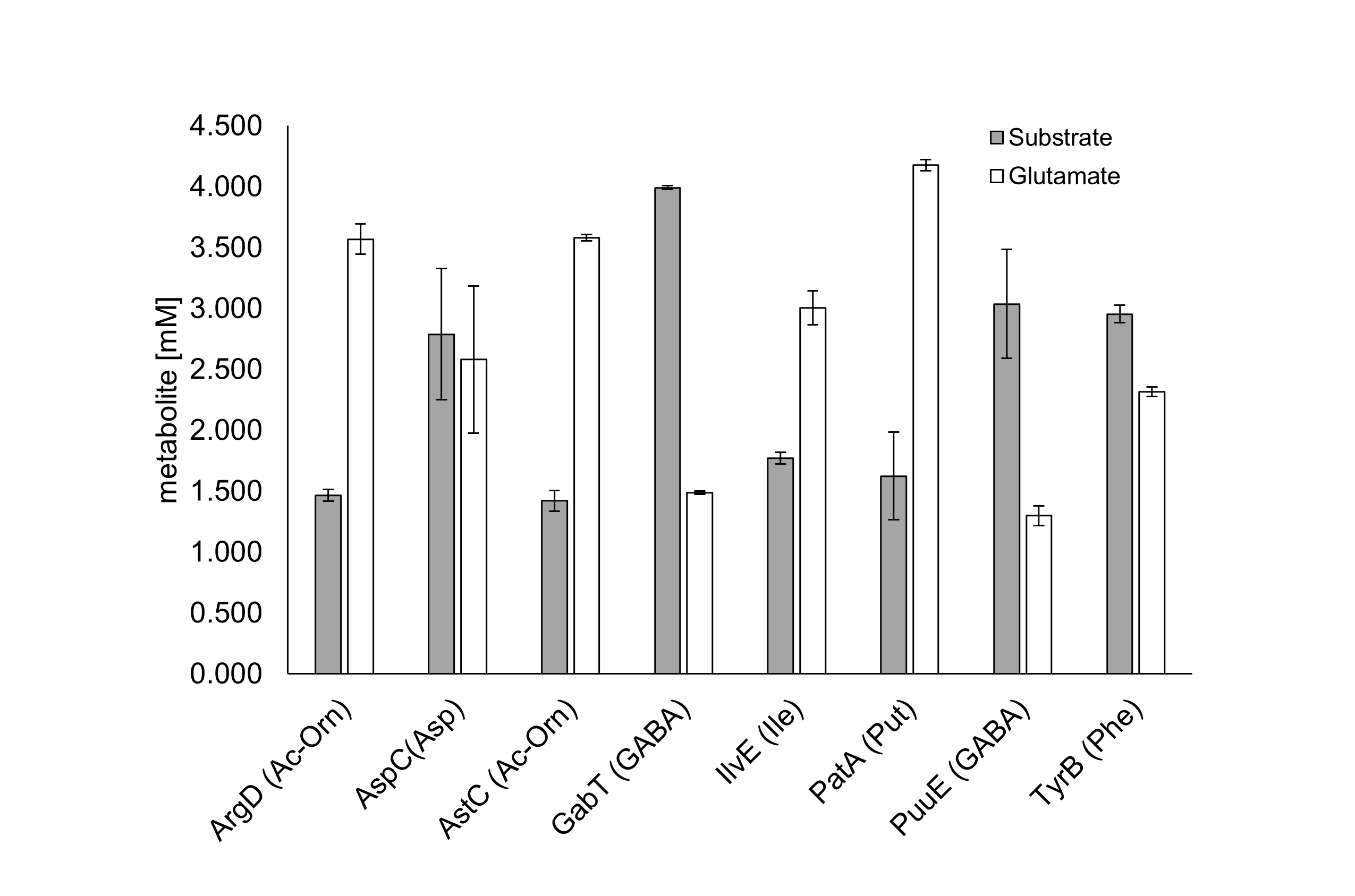

Figure S7. Production of glutamate by the purified transaminases used in this study, in presence of their native substrate. The concentration of glutamate was determined via HPLC-UV from reaction samples stopped after 30 min. Abbreviations: *N*_α_-Acetyl-ornithine (Ac-Orn), aspartate (Asp), γ-amino butyrate (GABA), isoleucine (Ile), putrescine (Put), phenylalanine (Phe). The data are shown as mean and SD of three independent replicates (n = 3). Dansylation and HPLC-UV analysis were performed as published previously (Aveta et al., 2026).

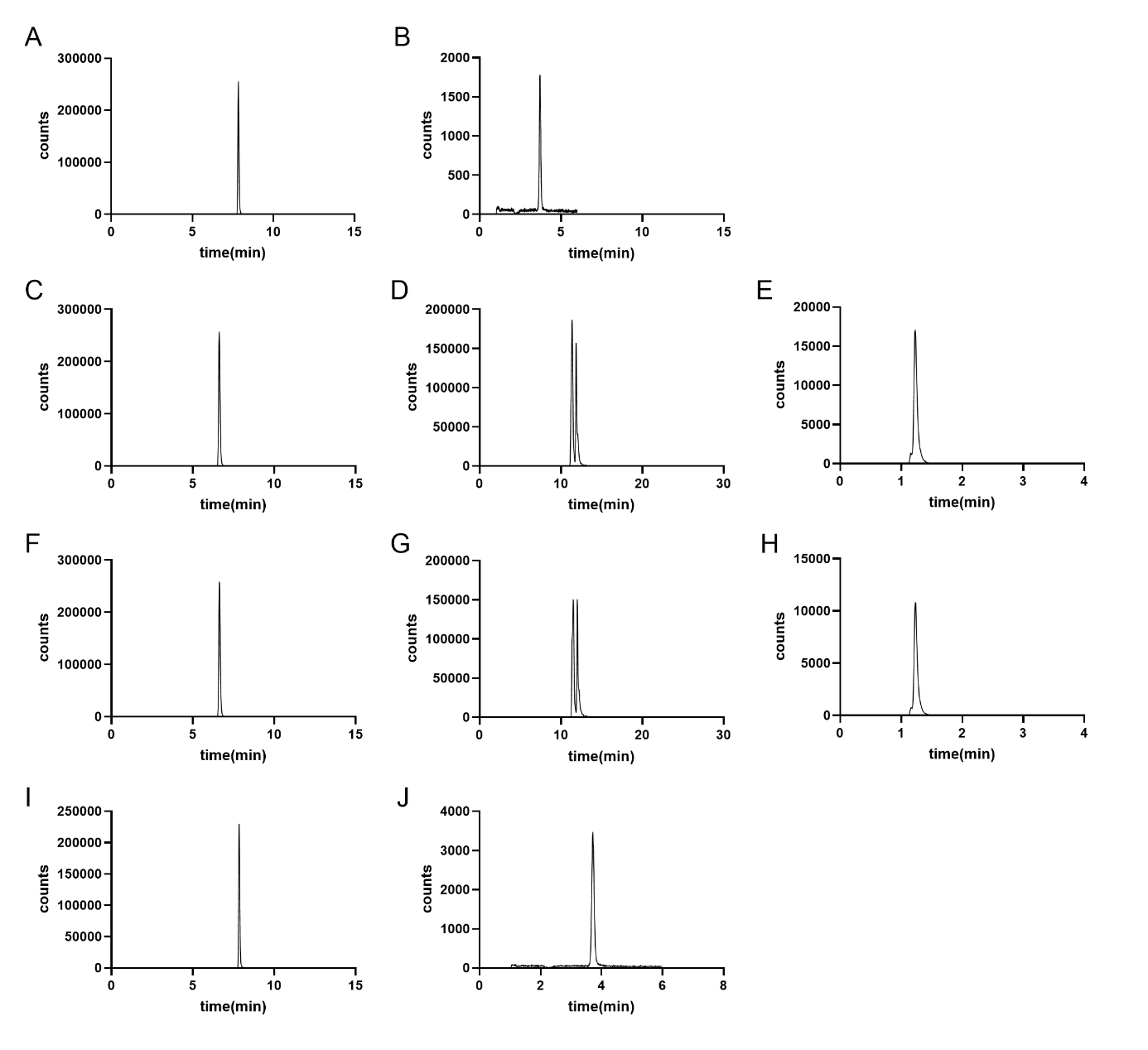

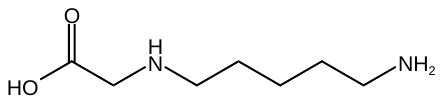

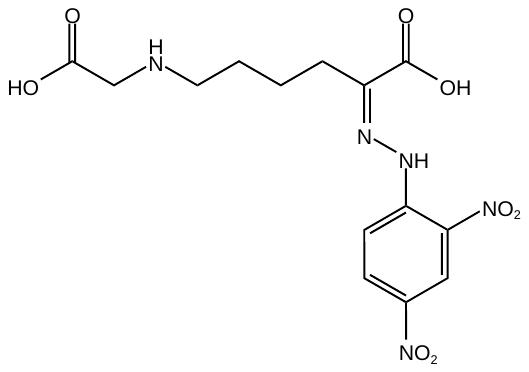

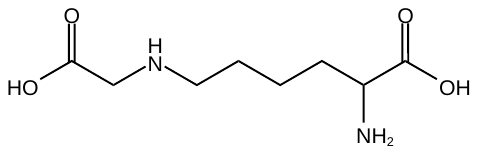

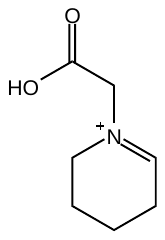

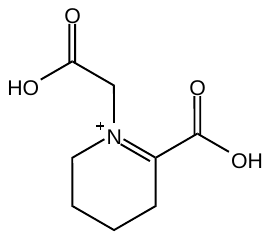

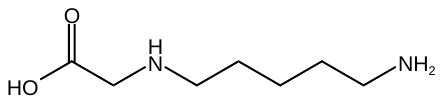

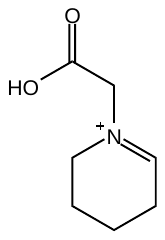

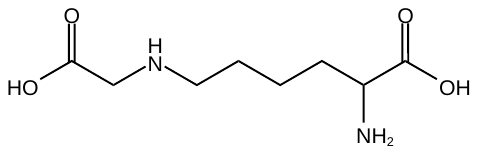

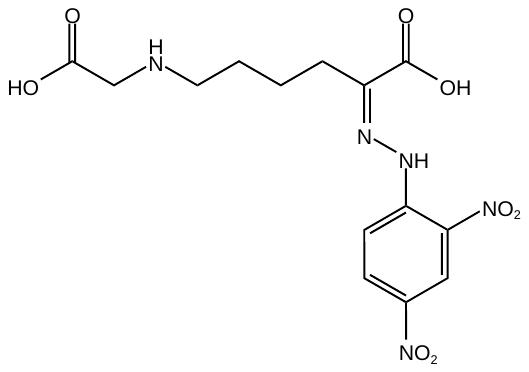

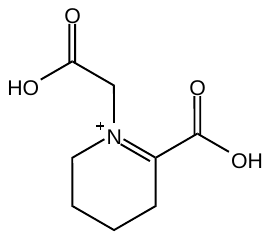

Figure S8. HPLC-MS of the *in vitro* reaction samples of GabT (first row), IlvE (second row), and PatA (third and fourth rows). The samples were collected after 24 h of incubation. A) the substrate CM-Cad in presence of GabT, and B), the resulting CM-piperideinium ion. C) The substrate CML in presence of IlvE, and the two products resulting D) CM-amino-oxo-hexanoic acid and E) CM-THPA. F) the substrate CML in presence of PatA and the related products G) CM-amino-oxo-hexanoic acid, and H) CM-THPA. I) The substrate CM-Cad in presence of PatA and the resulting product J) CM-piperideinium ion. The product CM-aminooxohexanoic acid was detectable through derivatization with 2,4-dinitrophenylhydrazine (DNPH), and the double peak is due to the formation of isomeric hydrazones.

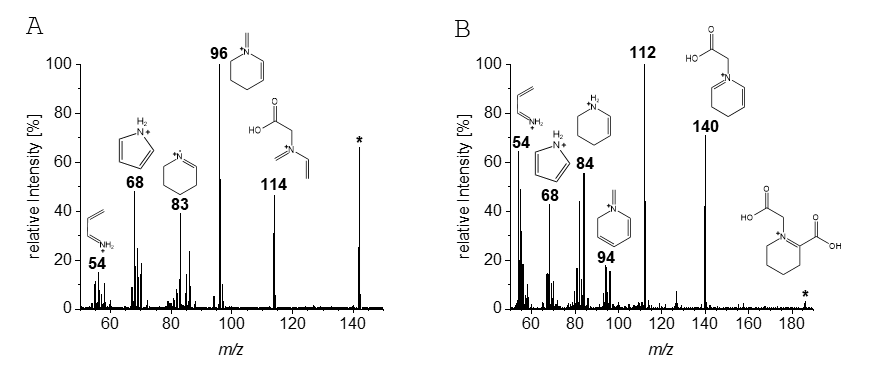

Figure S9. Postulated fragments (Grossert & White, 2021; Hellwig et al., 2019) of the identified products A) CM-piperideinium ion and B) CM-THPA. The two MS/MS spectra obtained by product ion scans (20 eV collision energy; 80 V fragmentor voltage) shown belong to the PatA samples containing CM-Cad and CML as substrates, respectively.

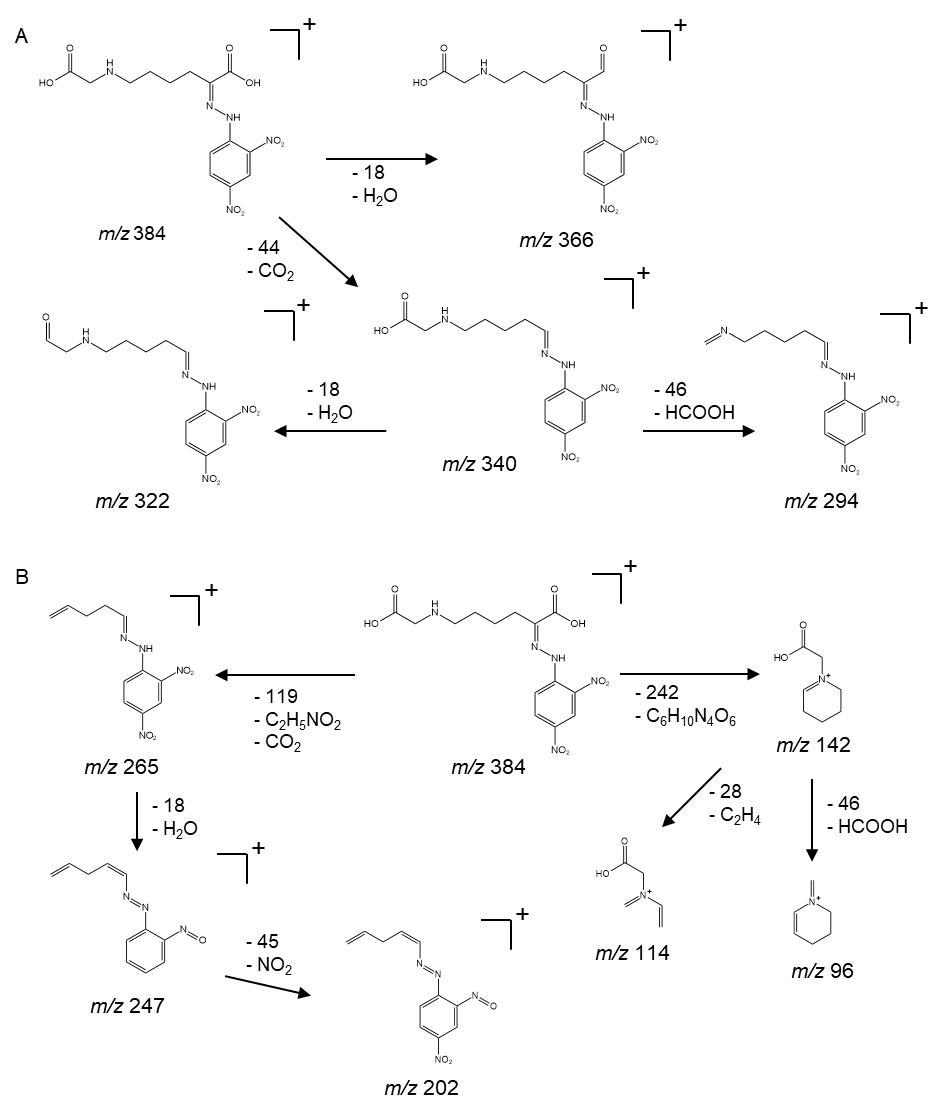

Figure S10. Postulated fragmentation pattern of the dinitrophenylhydrazone (Benoit & Holmes, 2011; Dator et al., 2016; Hellwig et al., 2019) of CM-aminooxohexanoic acid based on the MS/MS spectra obtained for IlvE and PatA in presence of CML.

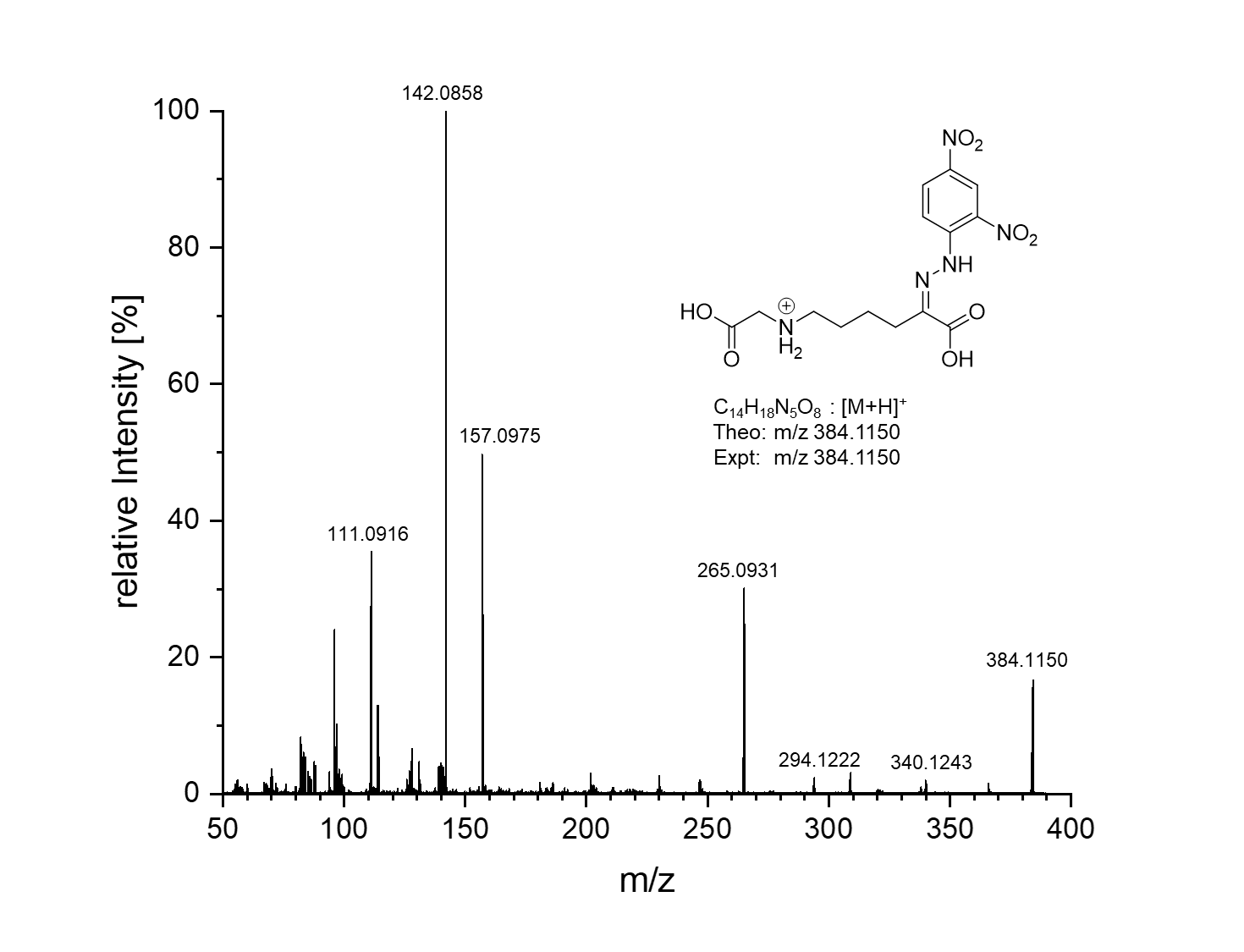

Figure S11. MS/MS spectrum of the precursor ion at *m/z* 384.1150 obtained via HPLC-MS/MS of a sample containing PatA and CML after 24 h of incubation
